# Host cell identity shapes the replication and evolution of *Drosophila* C virus

**DOI:** 10.64898/2026.08.02.742389

**Authors:** Qinyi Liang, Jiaxin Liu, Yunfan Huang, Xuye Yuan, Tatsuhiko Kadowaki

**Author notes:** Correspondence Tatsuhiko Kadowaki, PhD, Department of Biosciences and Bioinformatics, School of Science Xi’an Jiaotong-Liverpool University, 111 Ren’ai Road, Suzhou Dushu Lake Higher Education Town Jiangsu Province 215123, China.

## Abstract

RNA viruses encounter diverse cellular environments within their hosts, yet how different host cell types influence viral evolution remains poorly understood. Here, we investigated the replication and experimental evolution of Drosophila C virus (DCV) in glial, mesodermal, and hemocyte-derived S2 cells. Mesodermal cells supported substantially higher viral RNA accumulation, viral protein production, and infectivity than glial or S2 cells, demonstrating pronounced cell type-dependent permissiveness. Comparative transcriptomic and proteomic analyses revealed distinct host responses to DCV infection despite the common mesodermal origin of mesodermal and S2 cells. Functional analyses showed that RNA interference, cGAS–STING signaling, viral binding and entry, and cellular ATP abundance could not account for the enhanced replication of DCV in mesodermal cells, whereas S2 cells displayed stronger cGAS–STING and JAK– STAT responses. Serial passage of DCV for 15 generations in mesodermal or S2 cells resulted in rapid phenotypic adaptation, with evolved viruses exhibiting improved replication in the corresponding cell type without altering infectivity in adult flies. Genome sequencing revealed distinct evolutionary patterns between nonstructural and structural protein-coding regions. Nonstructural proteins remained highly conserved with limited evidence of parallel evolution, whereas structural proteins accumulated more mutations and exhibited broader signatures of positive selection. A recurrent VP3 mutation (G8089A; R270H) independently emerged in three S2-passaged lineages, representing the only mutation consistently associated with adaptation to a specific cell type. Structural modeling suggested that this substitution is unlikely to substantially alter VP2–VP3 interactions, implying more subtle effects on capsid function. Together, our findings demonstrate that different *Drosophila* cell types impose distinct intracellular selective pressures that shape DCV replication and evolutionary trajectories, highlighting host cell identity as an important determinant of RNA virus evolution.

## Introduction

RNA viruses evolve rapidly because of their high mutation rates, large population sizes, and short generation times. This remarkable evolutionary capacity enables viruses to adapt to changing host environments and immune or pharmacological pressures and, in some cases, expand their host range [1, 2]. Although virus evolution is often considered at the level of whole organisms or populations, viruses replicate within individual cells that differ substantially in their physiology, metabolism, and immune status. Consequently, different cell types may provide distinct intracellular environments that influence viral replication and the selective pressures acting on viral populations. However, the contribution of host cell identity to RNA virus evolution remains incompletely understood.

Drosophila C virus (DCV), a positive-sense single-stranded RNA virus belonging to the family Dicistroviridae, is a common and well-characterized natural viral pathogen of *Drosophila melanogaster* [3, 4]. DCV displays broad, route-dependent tissue tropism, including the digestive tract, fat body, visceral and somatic muscles, trachea, and reproductive tissues, resulting in tissue-specific pathology and host responses [5, 6]. As with other RNA viruses, successful DCV infection depends on extensive interactions with host cellular machinery, whereas host antiviral defenses restrict viral replication. In *Drosophila*, antiviral defense is mediated primarily by RNA interference (RNAi), with important virus-specific contributions from cGLR–STING and JAK–STAT signaling [7–11]. Genome-wide genetic and functional studies have identified host factors involved in viral entry, IRES-dependent translation, membrane trafficking, and fatty acid biosynthesis that are required for efficient DCV replication [12–14]. Moreover, tissue-specific differences in antiviral signaling have recently been demonstrated in *Drosophila*, including differential regulation of STING-dependent responses [15]. These observations suggest that the intracellular environment encountered by DCV varies substantially among different host cell types.

Experimental evolution has become a powerful approach for investigating virus adaptation under defined selective conditions. Serial-passage experiments have revealed how RNA viruses adapt to new hosts, cultured cells, antiviral compounds, and immune pressures, whereas deep sequencing enables the genetic dynamics underlying these adaptations to be followed at high resolution [16, 17]. Although many experimental-evolution studies use a single cell line, direct comparisons of viral evolution across distinct cellular environments remain relatively limited. Studies of dengue and Zika viruses have nevertheless demonstrated that propagation in different cell types can produce distinct phenotypic and genetic evolutionary outcomes [18, 19]. Whether different cell types originating from the same host species impose distinct selective pressures that drive divergent viral evolutionary trajectories remains poorly understood.

In this study, we investigated the replication and experimental evolution of DCV in three *Drosophila* cell types: glial, mesodermal, and hemocyte-like S2 cells. Continuous glial and mesodermal cell lines were generated by lineage-restricted expression of activated Ras, Ras^V12^ [20]. We first compared viral replication, host transcriptomic and proteomic responses, and the activities of major antiviral pathways to identify cellular factors associated with differential permissiveness to DCV infection. We then serially passaged DCV independently in mesodermal and S2 cells and analyzed the resulting viral populations using phenotypic assays and whole-genome sequencing. Our results demonstrate that different *Drosophila* cell types provide distinct intracellular environments that influence both viral replication and evolutionary trajectories, highlighting host cell identity as an important determinant of RNA virus evolution. Coleman-Gosser and colleagues directly describe the generation and characterization of the glial and mesodermal lines used in this type of work.

## Results

### DCV exhibits distinct cell tropism in *Drosophila* cell lines

To establish a cell culture system for experimental evolution, we first compared the susceptibility of three *Drosophila* cell lines—glial, mesodermal, and S2 cells— to DCV infection. Equal numbers of each cell type (Fig. 1A) were infected with the same dose of DCV (5 × 10^7^ TCID50), and viral replication was monitored at 6, 12, and 24 h post-infection by RNA-seq and quantitative proteomic analyses. Both viral genomic RNAs (ORF1 and ORF2) increased progressively in all three cell lines, indicating productive infection; however, viral RNA accumulated to significantly higher levels in mesodermal cells than in glial or S2 cells, particularly at 24 h post-infection (Fig. 1B). Consistent with the transcriptomic data, the abundance of the ORF1 and ORF2 polyproteins was also highest in mesodermal cells throughout the infection time course (Fig. 1C). Independent validation by RT-qPCR and immunoblotting further confirmed significantly higher levels of viral genomic RNA and the capsid protein VP1 in mesodermal cells than in glial or S2 cells at 24 h post-infection (Fig. 1D–F). Together, these results demonstrate that DCV displays marked cell type-dependent tropism in *Drosophila* cell lines, with mesodermal cells supporting substantially more efficient viral replication than glial or S2 cells.

**Figure 1.**
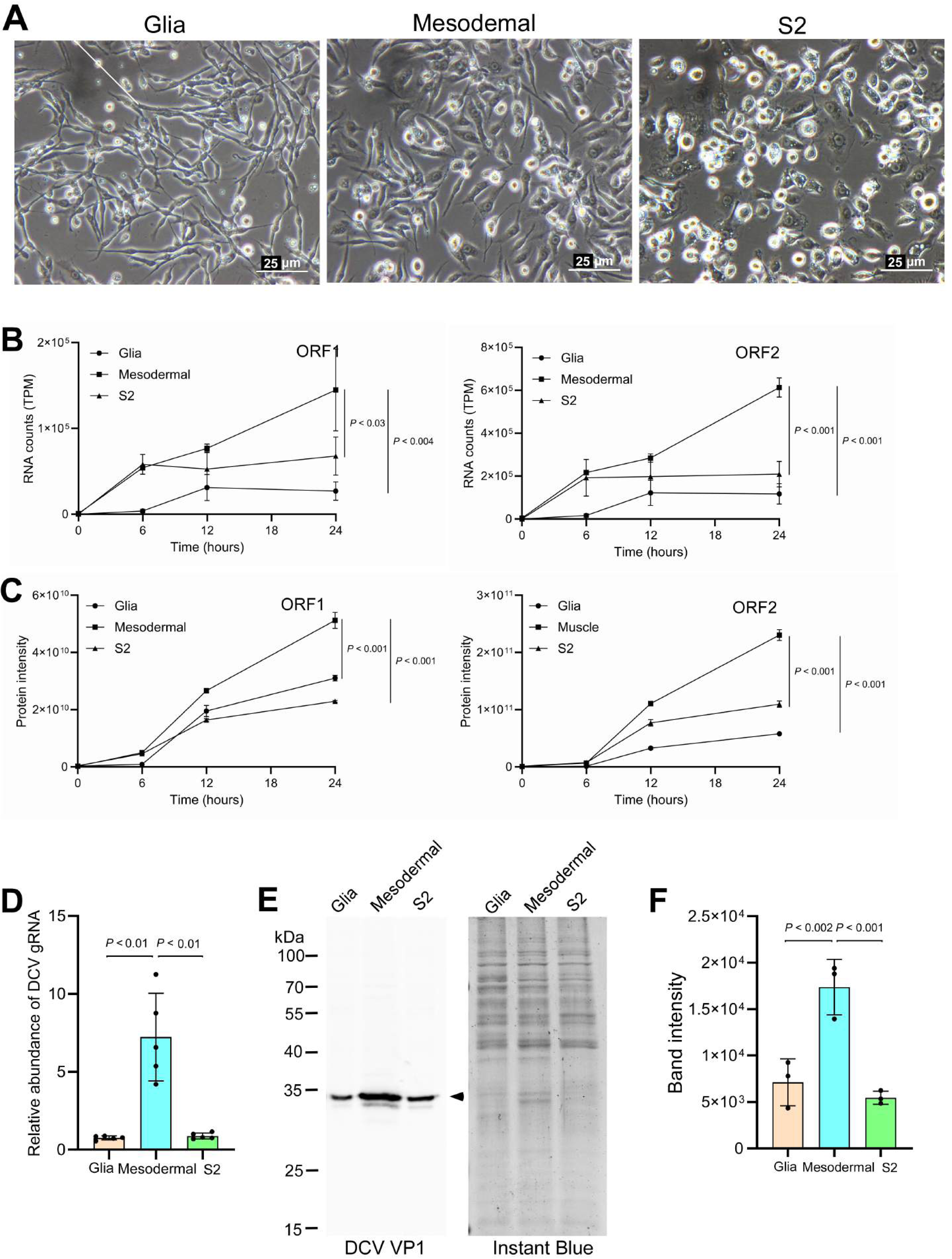
Mesodermal cells are highly permissive to DCV infection. (A) Morphology of glial, mesodermal, and S2 cells. Scale bar, 25 μm. (B) Abundance of DCV genomic RNA derived from ORF1 and ORF2 in the transcriptomes of glial, mesodermal, and S2 cells at 6, 12, and 24 h post-infection (hpi). At 24 hpi, mesodermal cells contained significantly more ORF1 RNA than glial (*P* < 0.004) or S2 cells (*P* < 0.03), and significantly more ORF2 RNA than either glial or S2 cells (*P* < 0.001). Statistical significance was assessed by one-way ANOVA followed by one-sided Dunnett’s multiple-comparison test; n = 3 biological replicates. (C) Abundance of DCV-derived peptides encoded by ORF1 and ORF2 in the proteomes of glial, mesodermal, and S2 cells at 6, 12, and 24 hpi. At 24 hpi, mesodermal cells contained significantly more ORF1- and ORF2-derived peptides than glial or S2 cells (*P* < 0.001 for each comparison). Statistical significance was assessed by one-way ANOVA followed by one-sided Dunnett’s multiple-comparison test; n = 3 biological replicates. (D) Relative abundance of DCV genomic RNA (gRNA) in glial, mesodermal, and S2 cells at 24 hpi. DCV gRNA abundance was highest in mesodermal cells. Statistical significance was assessed using the Kruskal–Wallis test followed by one-sided Steel multiple-comparison tests; n = 5 biological replicates. (E) Immunoblot analysis of DCV VP1 in lysates of infected glial, mesodermal, and S2 cells at 24 hpi. Total protein visualized by InstantBlue staining served as the loading control. The VP1 band is indicated by an arrowhead, and molecular mass markers are shown in kilodaltons (kDa) on the left. (F) Quantification of VP1 abundance in panel E. VP1 levels were significantly higher in mesodermal cells than in glial or S2 cells. Statistical significance was assessed by one-way ANOVA followed by one-sided Dunnett’s multiple-comparison test; n = 3 biological replicates.

### DCV infection elicits distinct transcriptomic and proteomic responses in glial, mesodermal, and S2 cells

To define host responses associated with the different cellular outcomes of DCV infection, we characterized the transcriptomes and proteomes of infected glial, mesodermal, and S2 cells. Principal component analysis (PCA) of the transcriptomic data showed that mesodermal and S2 cells clustered together and were clearly separated from glial cells (Fig. 2A), consistent with the mesodermal origin of hemocyte-like S2 cells. In contrast, proteomic profiles formed three distinct clusters corresponding to each cell type (Fig. 2B), indicating that cell type-specific differences were more pronounced at the protein level than at the transcript level. Although a small number of transcriptomic outliers were observed, the overall clustering pattern remained robust.

**Figure 2.**
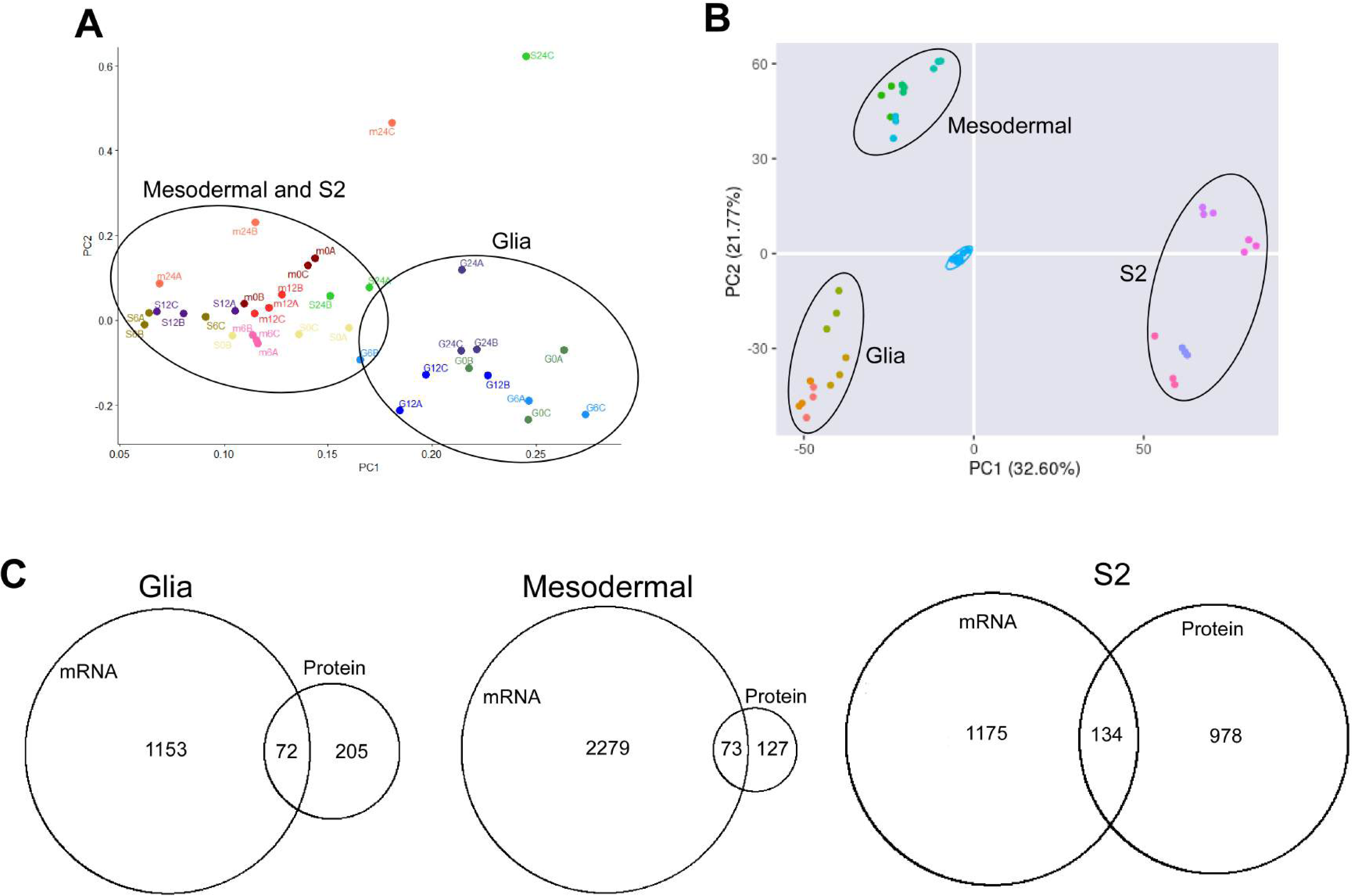
Transcriptomic and proteomic responses of glial, mesodermal, and S2 cells to DCV infection. (A) Principal component analysis (PCA) of transcriptomes from uninfected and DCV-infected glial, mesodermal, and S2 cells collected at 6, 12, and 24 h post-infection. Transcriptomic profiles of mesodermal and S2 cells clustered together and were separated from those of glial cells. Two samples, one mesodermal and one S2 sample at 24 h post-infection, were identified as outliers. (B) PCA of proteomes from uninfected and DCV-infected glial, mesodermal, and S2 cells collected at 6, 12, and 24 h post-infection. Proteomic profiles formed distinct clusters according to cell type. Quality-control samples are shown in light blue. (C) Venn diagrams showing the numbers and overlap of mRNAs and proteins significantly altered by DCV infection in glial, mesodermal, and S2 cells.

DCV infection induced the greatest transcriptional response in mesodermal cells, whereas the largest changes in protein abundance occurred in S2 cells (Fig. 2C). Moreover, transcriptomic changes substantially exceeded proteomic changes in glial and mesodermal cells, whereas the numbers of differentially expressed mRNAs and proteins were comparable in S2 cells (Fig. 2C). These results suggest that host responses to DCV infection are regulated differently among the three cell types and that post-transcriptional regulation may contribute more extensively to the antiviral response in S2 cells.

To compare the biological processes altered by DCV infection, differentially expressed mRNAs and proteins were integrated into STRING interaction networks and clustered into functional pathway modules (Supplementary Data 1). Several modules, including ribosome and translation, microtubule organization, chromatin organization, mitochondrial function, lipid and fatty acid metabolism, RNA splicing, DNA repair, and glutathione transferase activity, were shared among all three cell types, indicating a conserved cellular response to DCV infection. In contrast, autophagy, actin filament organization, and protein folding modules were shared only between glial and S2 cells, whereas additional cell type-specific modules were identified in each cell line. Glial cells uniquely exhibited enrichment of the chemical synaptic transmission module, whereas mesodermal cells showed specific enrichment of nucleocytoplasmic transport and regulation of cell death modules, with most associated genes being upregulated following infection except Drice and Diap1.

Notably, S2 cells uniquely displayed enrichment of immune response, JAK–STAT signaling, and viral entry into host cell modules. Genes associated with immune response and JAK–STAT signaling were predominantly upregulated, whereas genes assigned to the viral entry module were largely downregulated. Together, these findings demonstrate that, despite their shared developmental origin, mesodermal and S2 cells mount distinct molecular responses to DCV infection, with S2 cells exhibiting a more prominent immune-related transcriptional program.

### Differences in DCV permissiveness cannot be explained by RNAi, cGAS– STING, or JAK–STAT signaling

The marked differences in DCV replication among glial, mesodermal, and S2 cells prompted us to investigate whether variation in the major antiviral pathways contributes to their differential permissiveness. We therefore compared the activities of the RNA interference (RNAi), cGLR–STING, and JAK–STAT pathways in the three cell types.

To evaluate RNAi activity, glial, mesodermal, and S2 cells were co-transfected with a GFP expression plasmid and either GFP-specific or control (mCherry) dsRNA. GFP fluorescence was readily detected in cells transfected with control dsRNA, although the overall fluorescence intensity varied among the three cell types, likely reflecting differences in transfection efficiency (Fig. 3A). Co-transfection with GFP dsRNA markedly reduced GFP fluorescence in all three cell types, as confirmed by fluorescence microscopy, flow cytometry, and immunoblotting (Fig. 3A–C). These results demonstrate that the RNAi pathway is functional in glial, mesodermal, and S2 cells.

**Figure 3.**
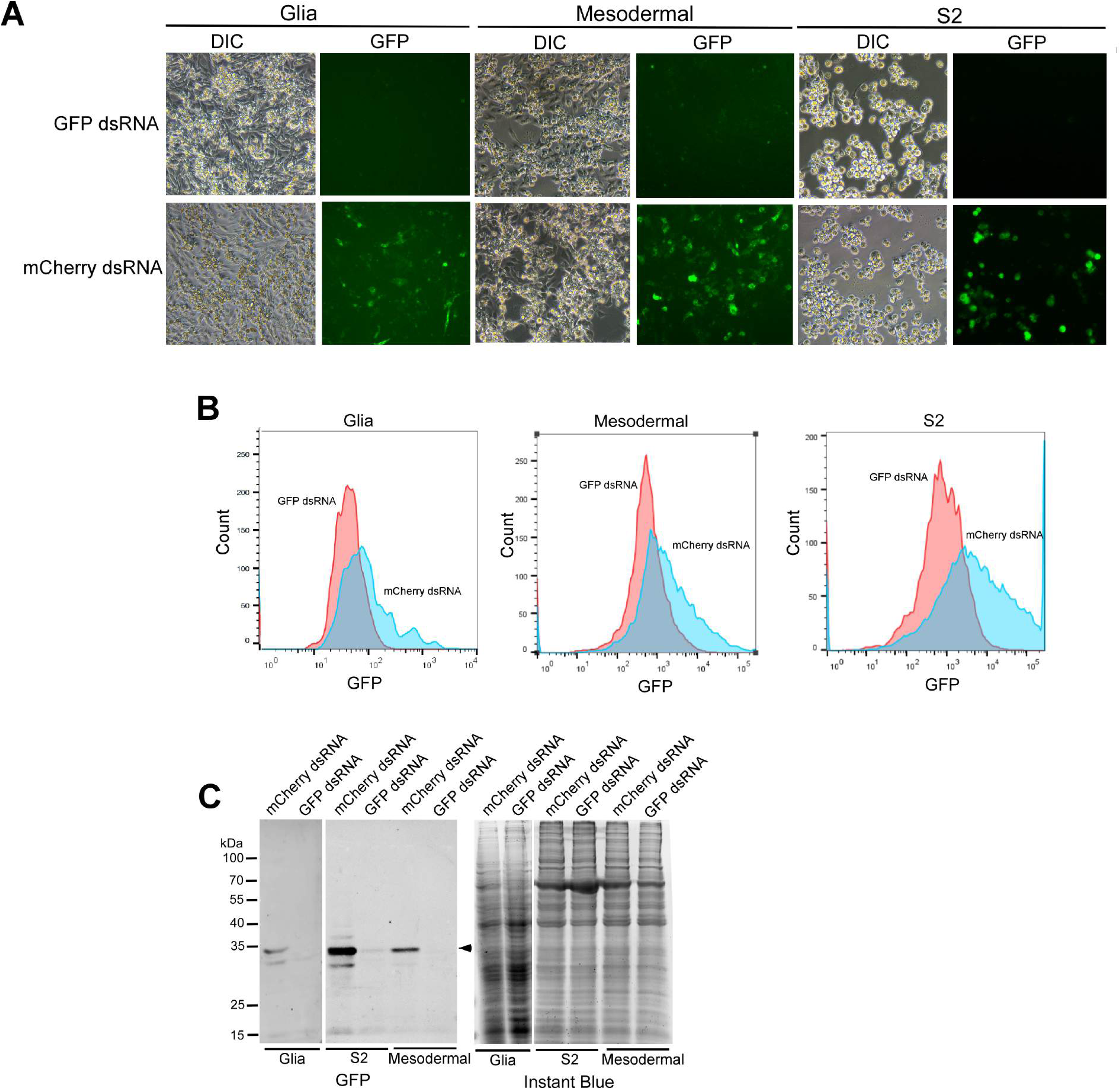
RNA interference is functional in glial, mesodermal, and S2 cells. (A) Differential interference contrast (DIC) and GFP fluorescence images of glial, mesodermal, and S2 cells co-transfected with a GFP expression plasmid and either GFP-specific or mCherry-specific control dsRNA. GFP exposure times were 600, 300, and 100 ms for glial, mesodermal, and S2 cells, respectively. (B) Flow cytometric analysis of GFP fluorescence in the cells shown in panel A. (C) Immunoblot analysis of GFP in lysates of the cells shown in panel A. Total protein visualized by InstantBlue staining served as the loading control. The GFP band is indicated by an arrowhead, and molecular mass markers are shown in kilodaltons (kDa) on the left. Co-transfection with GFP-specific, but not mCherry-specific, dsRNA reduced GFP expression in all three cell types.

We next examined cGLR–STING signaling by stimulating cells with 2′3′-cGAMP and measuring the expression of the downstream target gene *Srg3*. Treatment with 2′3′-cGAMP significantly induced *Srg3* expression in all three cell types, indicating that STING signaling is functional (Fig. 4A). However, both basal and cGAMP-induced *Srg3* expression were highest in S2 cells and lowest in mesodermal cells. Consistent with this observation, expression of *cGLR1*, an upstream component of the pathway, was significantly higher in S2 cells than in glial or mesodermal cells, which exhibited similarly low expression levels (Fig. 4B). These findings suggest that cGLR–STING signaling is less active in mesodermal cells than in S2 cells.

**Figure 4.**
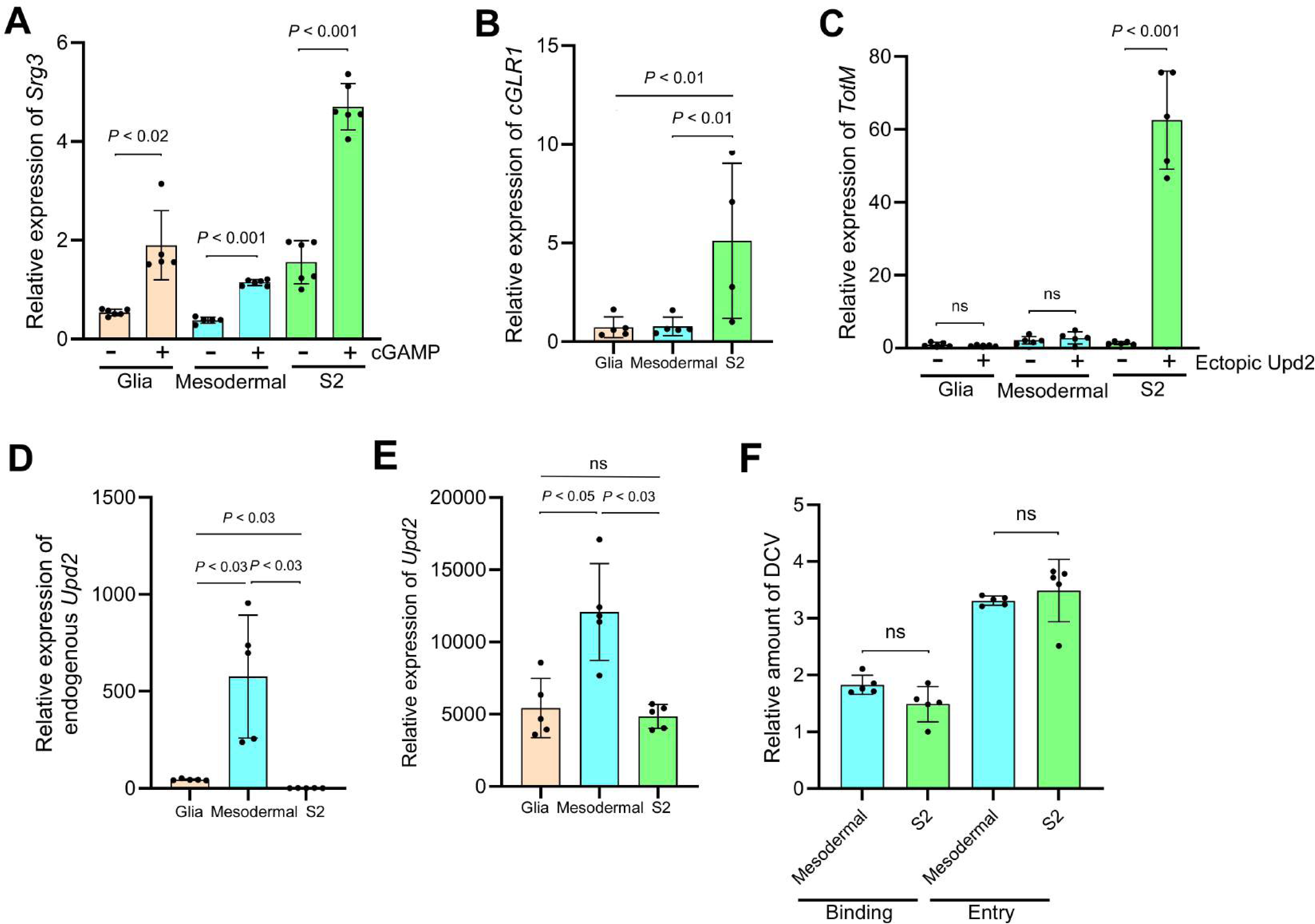
Comparison of cGAS–STING and JAK–STAT signaling as well as DCV binding and entry in glial, mesodermal, and S2 cells. (A) Relative expression of the STING-responsive gene *Srg3* in glial, mesodermal, and S2 cells treated with 2′3′-cGAMP (+) or vehicle control (−) for 8 h. Differences between treated and control cells were analyzed using two-sided Welch’s *t*-tests and comparisons among the three cell types were performed using the Kruskal– Wallis test followed by one-sided Steel multiple-comparison tests; *n* = 5-6 biological replicates. (B) Relative endogenous expression of *cGLR1* in glial, mesodermal, and S2 cells. Statistical significance was assessed using the Kruskal–Wallis test followed by one-sided Steel multiple-comparison tests; n = 4-5 biological replicates. (C) Relative expression of the JAK–STAT target gene *TotM* in cells transfected with an Upd2 expression plasmid (+) or empty vector (−). Differences between Upd2-expressing and control cells were analyzed using two-sided Welch’s *t*-tests and comparisons among the three cell types were performed using the Kruskal–Wallis test followed by the Steel–Dwass test; n = 5 biological replicates. (D) Relative endogenous expression of *Upd2* in glial, mesodermal, and S2 cells. (E) Relative *Upd2* expression following transfection with the Upd2 expression plasmid. For D and E, statistical significance was assessed using the Kruskal–Wallis test followed by Steel–Dwass multiple-comparison tests; n = 5 biological replicates. (F) Relative amounts of cell-associated DCV genomic RNA following viral binding at 4 °C for 30 min or entry at 25 °C for 2 h in mesodermal and S2 cells. Differences between the two cell types were analyzed using two-sided Welch’s *t*-tests; n = 5 biological replicates. Relative transcript and viral RNA levels were determined by RT–qPCR. Bars represent means, error bars indicate standard deviations, and dots represent biological replicates. ns, not significant.

Finally, we compared JAK–STAT signaling by ectopically expressing the cytokine Upd2 and measuring induction of the downstream target gene *TotM*. Ectopic *Upd2* expression induced approximately 50-fold upregulation of *TotM* in S2 cells but failed to activate *TotM* expression in either glial or mesodermal cells (Fig. 4C). Interestingly, glial and mesodermal cells expressed substantially higher levels of endogenous *Upd2* than S2 cells, yet basal *TotM* expression remained comparable among the three cell types (Fig. 4D). Following ectopic *Upd2* expression, *Upd2* transcript levels remained highest in mesodermal cells (Fig. 4E), indicating that the lack of *TotM* induction in glial and mesodermal cells is unlikely to result from insufficient ligand production. Up-regulation of *Upd2* in glia and mesodermal cells could be induced by activated Ras to establish the cell lines [20]. Together, these results indicate that JAK–STAT signaling is responsive to *Upd2* in S2 cells but is largely refractory in glial and mesodermal cells.

Collectively, these findings demonstrate that the enhanced permissiveness of mesodermal cells to DCV infection cannot be explained by defects in the RNAi pathway or by broad impairment of cGLR–STING or JAK– STAT signaling, suggesting that additional cell type-specific mechanisms underlie the distinct susceptibility of these cells to DCV infection.

### Enhanced DCV replication in mesodermal cells is independent of viral entry and cellular ATP abundance

The higher level of DCV replication in mesodermal cells could potentially result from more efficient viral attachment or entry. To test this possibility, we compared the efficiency of DCV binding and entry between mesodermal and S2 cells. No significant differences were detected in either viral binding or entry, indicating that the enhanced permissiveness of mesodermal cells is unlikely to result from increased viral attachment or internalization (Fig. 4F).

Because viral replication depends heavily on host energy metabolism, we next examined whether mesodermal cells contain higher intracellular ATP levels than S2 cells. Surprisingly, the estimated ATP concentration was higher in S2 cells (19.6 mM) than in mesodermal cells (8.37 mM). Although the absolute ATP concentrations may be overestimated because of limitations of the luciferase-based assay and cell volume estimation, these measurements clearly indicate that mesodermal cells do not possess a higher ATP concentration than S2 cells. Together, these findings suggest that neither differences in viral entry nor intracellular ATP abundance account for the enhanced replication of DCV in mesodermal cells.

### Serial passage drives cell type-dependent adaptation of DCV

Having established that mesodermal and S2 cells differ markedly in their permissiveness to DCV infection, we next asked whether these distinct intracellular environments impose different selective pressures on viral evolution. To minimize pre-existing viral diversity, the parental DCV population was first subjected to three rounds of bottlenecking by endpoint dilution in S2 cells. This bottlenecked virus was then serially passaged independently in mesodermal and S2 cells for 15 passages using identical inocula (10,000 TCID50) at each passage.

Viral infectivity increased rapidly during the initial passages in both cell types. The TCID50 values of viruses recovered from both mesodermal and S2 cells were higher at passage 5 than those of the parental bottlenecked virus (Fig. 5A and B), indicating early adaptation during serial passage. Thereafter, viral infectivity remained largely stable, although viruses propagated in S2 cells exhibited a further increase in infectivity between passages 5 and 10 when titrated on S2 cells, but not when titrated on mesodermal cells (Fig. 5A and B). Moreover, viruses passaged in mesodermal cells consistently displayed higher infectivity when titrated on mesodermal cells than on S2 cells, whereas viruses passaged in S2 cells showed similar infectivity in the two cell types except at passage 10 (Fig. 5C). Because the parental virus also exhibited substantially higher infectivity in mesodermal cells than in S2 cells (TCID50: 2.7 × 10^8^/mL vs 4.7 × 10^7^/mL), serial passage in S2 cells appears to shift this initial phenotype toward improved replication in S2 cells.

**Figure 5.**
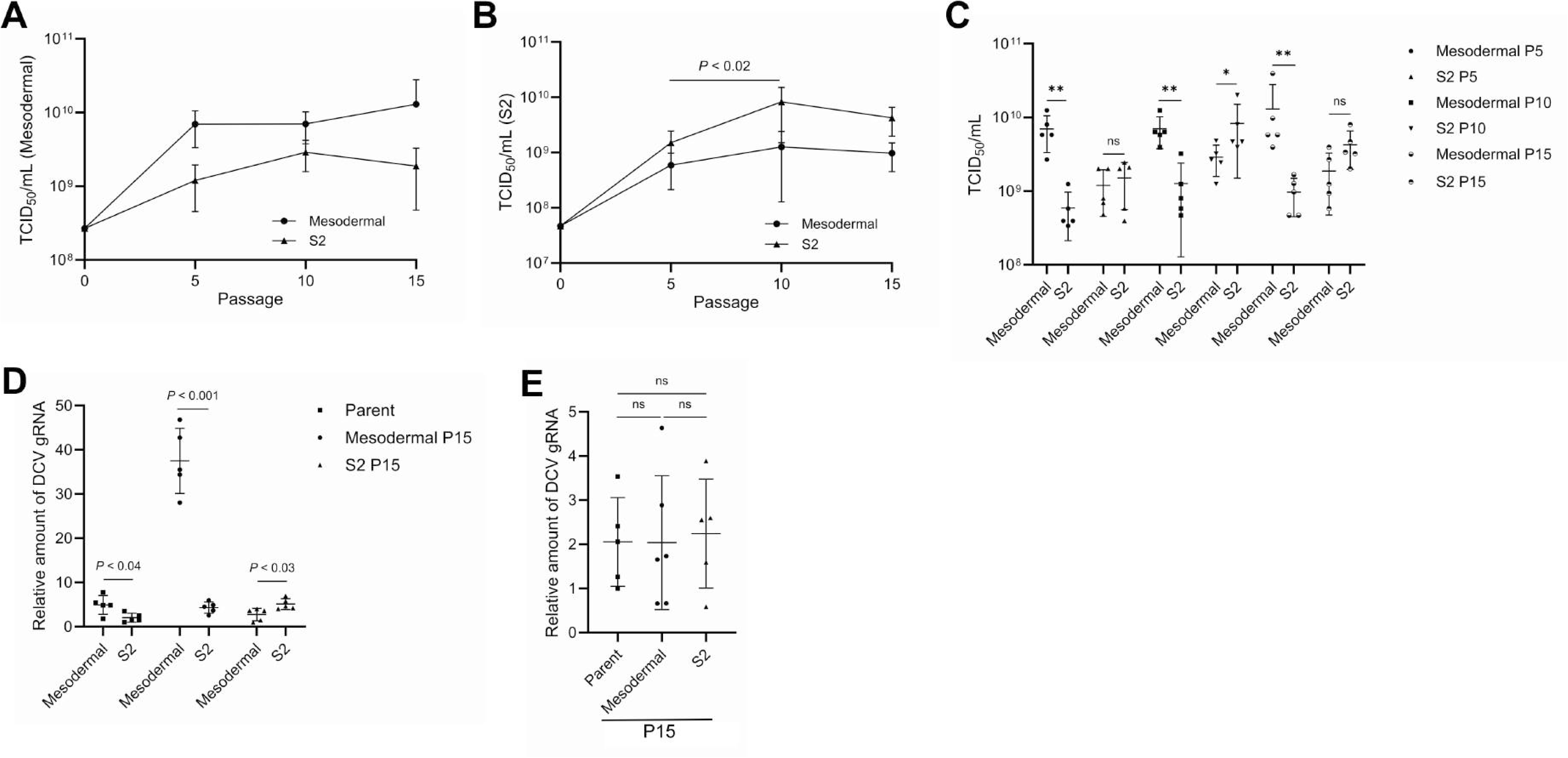
Experimental evolution of DCV in mesodermal and S2 cells. (A, B) Infectious titers of the parental DCV stock and virus populations serially passaged in mesodermal or S2 cells for 5, 10, and 15 passages. TCID50 values were determined using mesodermal cells (A) or S2 cells (B). Each point represents the mean of five independent viral lineages, and error bars indicate standard deviations. The titer of S2-passaged DCV increased significantly between P5 and P10 when measured in S2 cells (Kruskal–Wallis test followed by the Steel–Dwass test). (C) Comparison of the TCID50 values of mesodermal- and S2-passaged DCV populations at P5, P10, and P15 when titrated using mesodermal or S2 cells. Each symbol represents an independent viral lineage. Paired comparisons between titers determined in the two cell types were performed using Mann–Whitney *U* test. *, *P* < 0.05; **, *P* < 0.01; ns, not significant. (D) Relative abundance of DCV genomic RNA (gRNA) in mesodermal and S2 cells at 24 h after infection with the parental virus or pooled mesodermal P15 or S2 P15 viruses. Viral inputs were normalized according to gRNA abundance. Comparisons between mesodermal and S2 cells infected with the same viral stock were performed using two-sided Welch’s *t*-tests. Comparisons among the three viral stocks within each cell type were performed using the Kruskal–Wallis test followed by Steel–Dwass multiple-comparison tests; n = 5 biological replicates. (E) Relative DCV gRNA abundance in adult female *w*^1118^ flies at 24 h after intrathoracic injection with the parental virus or pooled mesodermal P15 or S2 P15 viruses. Each point represents a biological replicate consisting of ten flies; n = 5 biological replicates. Differences among the three viral populations were analyzed by one-way ANOVA. ns, not significant. In C-E, horizontal lines indicate means and error bars indicate standard deviations.

To determine whether serial passage altered viral replication in each cell type, mesodermal and S2 cells were infected with the parental virus or passage 15 (P15) viruses. To ensure equivalent viral input, inocula were normalized by viral genome copy number rather than TCID50, as the P15 virus stocks contained approximately twice the number of viral genomes as the parental stock. As expected, mesodermal cells accumulated higher levels of viral RNA than S2 cells following infection with the parental virus (Fig. 5D), consistent with the results shown in Fig. 1. However, viruses passaged in mesodermal cells produced the highest level of viral RNA in mesodermal cells, whereas viruses passaged in S2 cells produced higher levels of viral RNA in S2 cells than did the parental virus (Fig. 5D). Similar trends were observed for VP1 protein accumulation, although differences in S2 cells did not reach statistical significance (Supplementary Fig. 1). These findings indicate that independent passage in different cell types alters viral infectivity in a cell type-dependent manner.

Finally, we examined whether adaptation acquired during cell culture passage affected infection in adult flies. Despite the altered replication characteristics observed in cultured cells, parental, mesodermal P15, and S2 P15 viruses exhibited comparable infectivity in adult *Drosophila* (Fig. 5E), indicating that adaptation to either cultured cell type does not measurably alter viral infection in the whole-animal host.

### Serial passage reveals cell type-specific evolutionary trajectories of DCV

To identify mutations accumulated during serial passage in mesodermal and S2 cells, we first pooled DCV populations from five independent lineages at P5, P10, and P15 and sequenced the complete viral genomes using Illumina sequencing. Single nucleotide variations (SNVs) were distributed throughout the DCV genome in both cell types (Fig. 6A). However, three mutations (G6079T, G6082A, and T6181A) within the intergenic region (IGR) IRES reached high frequencies in both mesodermal- and S2-passaged viruses. Because T6079, A6082, and A6181 are the predominant nucleotides among published DCV genome sequences, whereas our parental virus contained G6079, G6082, and T6181, these mutations likely represent reversion of variants that became fixed during the bottlenecking process rather than adaptation to either cell type. Non-synonymous mutations were predominantly clustered within the genomic regions encoding 1A, 2B, 2C (helicase), and the structural proteins (Fig. 6B). We therefore sequenced these two regions separately in each of the three independent lineages. In contrast, only a few non-synonymous mutations were detected in the 3A and RNA-dependent RNA polymerase (RdRp) coding regions, and none were detected in the VPg or 3C protease coding regions, suggesting that these viral proteins are subject to strong purifying selection.

**Figure 6.**
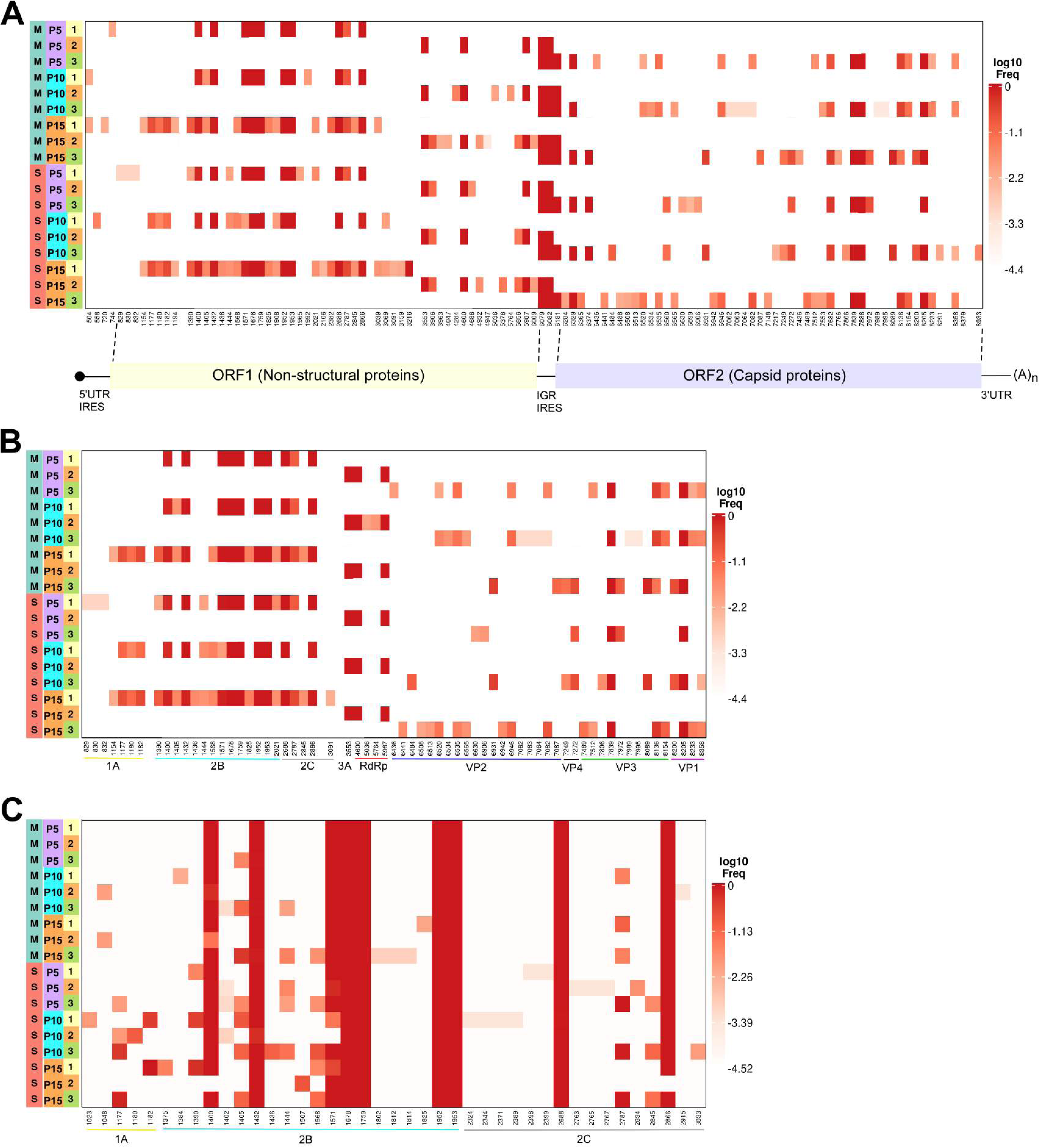
Genome-wide mutation profiles of DCV during serial passage in mesodermal and S2 cells. (A) Heatmap showing the frequencies of all single-nucleotide variants (SNVs) detected across the DCV genome in pooled virus populations passaged in mesodermal (M) or S2 (S) cells at passages 5, 10, and 15 (P5, P10, and P15). Virus populations from five independent lineages were pooled at each passage and amplified as three overlapping genomic fragments, indicated by 1–3. Genomic coordinates are based on the DCV EB reference genome. The genome organization, including the 5′-UTR IRES, ORF1 encoding nonstructural proteins, intergenic-region (IGR) IRES, ORF2 encoding capsid proteins, 3′ UTR, and poly(A) tail, is shown below the heatmap. (B) Heatmap showing the frequencies and genomic positions of nonsynonymous SNVs identified in the pooled virus populations shown in (A). The viral proteins encoded by the corresponding genomic regions are indicated below the heatmap. (C) Heatmap showing the frequencies and positions of nonsynonymous SNVs within the ORF1 region encoding 1A, 2B, and 2C in three independently evolved viral lineages at P5, P10, and P15. In this panel, 1–3 denote the three independent lineages. In all panels, color intensity represents the log10-transformed SNV frequency, and white indicates that the variant was not detected.

Within the ORF1 region encoding 1A, 2B, and 2C, the number of SNVs remained relatively constant during serial passage. The number of SNVs differed between mesodermal- and S2-passaged viruses only at P5 (Supplementary Fig. 2A). Most SNVs occurred at frequencies between 0.5 % and 50 % in both cell types, with no significant difference in their frequency distribution (Supplementary Fig. 2B). Mean nucleotide diversity differed between mesodermal- and S2-passaged viruses only at P10 (Supplementary Fig. 2C). Among the three ORF1 proteins, only 2B showed significantly higher nucleotide diversity than the mean diversity of the ORF1 region in S2 cells at P10 (Supplementary Fig. 2D). Consistent with this, πN–πS analysis detected positive selection acting only on the 2B gene in S2 cells at P10 and P15 (Supplementary Fig. 2E). Nevertheless, comparison of non-synonymous mutations across three independent lineages revealed no recurrent mesodermal- or S2-specific mutations in 1A, 2B, or 2C (Fig. 6C), indicating that parallel adaptation was not evident in the ORF1 region.

Within ORF2, which encodes the structural proteins, the number of polymorphic sites reached means of 24.0 and 20.3 in mesodermal- and S2-passaged populations, respectively, at P15; however, the increase was significant only in mesodermal-passaged viruses between P5 or P10 and P15 (Supplementary Fig. 3A). Most SNVs occurred at frequencies of 0.5–50 % in both cell types, with a higher proportion of intermediate-frequency SNVs in mesodermal-passaged populations and of high-frequency SNVs (>50 %) in S2-passaged populations (Supplementary Fig. 3B). Mean nucleotide diversity did not change significantly over time in either cell type (Supplementary Fig. 3C). Only VP3 showed significantly greater nucleotide diversity than the mean for the entire ORF2 region, and only in mesodermal cells at P5, indicating that no structural protein consistently accumulated more mutations during passage (Supplementary Fig. 3D). πN–πS analysis suggested positive selection on VP2 and VP3 at all passages in mesodermal cells, whereas in S2 cells it was detected for most structural proteins, except VP2 at P5 and VP4 and VP1 at P10 (Supplementary Fig. 3E). Thus, different structural protein-coding genes experienced positive selection in the two cell types.

Figure 7A shows the distribution of non-synonymous mutations in the ORF2 region across three independently evolved DCV lineages at P5, P10, and P15. The A8358G mutation was detected specifically in mesodermal-passaged viruses in multiple lineages, except lineage 1 at P10 and lineage 3 at P15. This mutation substitutes I62 of VP1 with valine but remained at a low frequency, reaching a maximum of only 6.4 %, suggesting that it is unlikely to represent an adaptive mutation. In contrast, the G8089A mutation in VP3 arose specifically in S2-passaged viruses, except in lineage 3 at P5, and was detected in all three independent lineages, indicating parallel evolution under S2 cell-specific selective pressure. The frequency of G8089A varied widely, ranging from 2.0 % (lineage 3 at P10) to 99.6 % (lineage 1 at P15), suggesting either relatively weak selection or that this site represents a mutational hotspot in S2 cells. G8089A results in an R270H substitution in VP3. Sequence alignment showed that this region of VP3 is poorly conserved among dicistroviruses, with an alignment gap immediately downstream of R270 (Fig. 7B). AlphaFold3 prediction of the DCV VP1–VP4 capsid complex placed R270 within a C-terminal α-helix of VP3, where its side chain is positioned near P144, N145, and S148 of VP2 but does not appear to contact them directly. Instead, R270 primarily interacts with residues within the VP3 α-helix (Fig. 7C). The corresponding residue in Cricket paralysis virus (CrPV), R273, occupies a similar structural position but is predicted to interact with T150 of VP2 (Fig. 7D). These structural analyses suggest that the R270H substitution is unlikely to substantially alter the interaction between VP3 and VP2.

**Figure 7.**
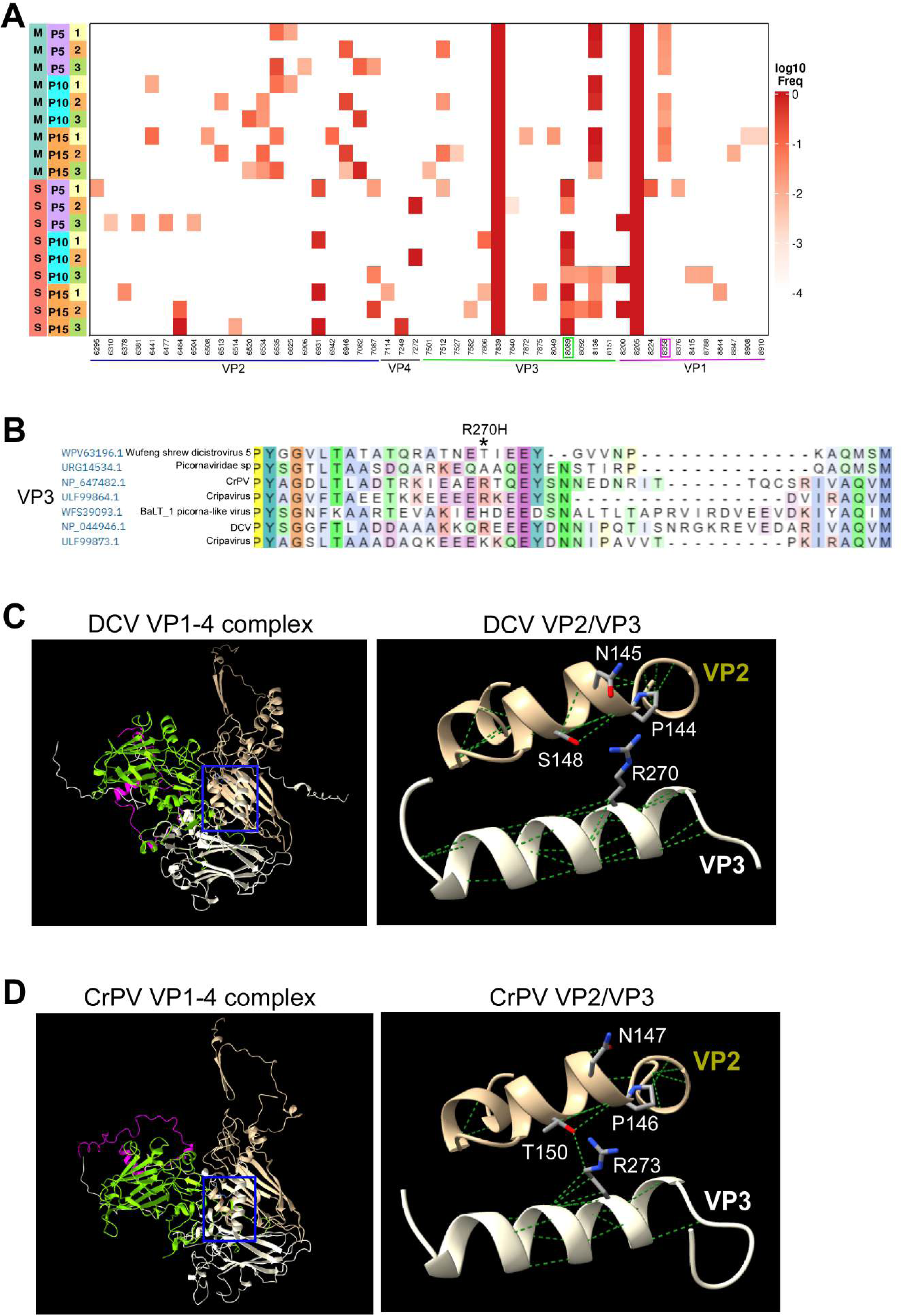
Recurrent emergence and structural context of the DCV VP3 R270H substitution. (A) Heatmap showing the frequencies and genomic positions of nonsynonymous SNVs within ORF2 in three independently evolved DCV lineages passaged in mesodermal (M) or S2 (S) cells at passages 5, 10, and 15 (P5, P10, and P15). The corresponding structural proteins are indicated below the heatmap. The S2-associated G8089A mutation in VP3 and the mesodermal-associated A8358G mutation in VP1 are outlined in green and magenta, respectively. Color intensity represents the log10-transformed SNV frequency, and white indicates that the variant was not detected. (B) Multiple-sequence alignment of the VP3 region surrounding DCV residue R270 among seven dicistroviruses. The R270H substitution is indicated above the alignment with asterisk. (C) AlphaFold3-predicted structure of the DCV VP1–VP4 complex and enlarged view of the VP2– VP3 interface indicated by the blue box. VP1-VP4 proteins are shown in green, khaki, light grey, and magenta, respectively. VP3 residue R270 and the nearby VP2 residues P144, N145, and S148 are shown as sticks. R270 is positioned near these VP2 residues but does not appear to contact them directly. Amino acid interactions are shown in green dashed lines. (D) Structure of the Cricket paralysis virus (CrPV) VP1–VP4 complex (PDB: 1B35) and enlarged view of the corresponding VP2–VP3 interface. The DCV R270-equivalent residue, R273 of CrPV VP3, and the nearby VP2 residues P146, N147, and T150 are shown as sticks. R273 interacts with T150 of VP2. Protein structures were visualized using UCSF ChimeraX.

## Discussion

Viruses encounter diverse cellular environments within their hosts, yet how these environments influence viral evolution remains poorly understood. In this study, we demonstrate that different *Drosophila* cell types support distinct levels of DCV replication and impose different evolutionary pressures during serial passage. Mesodermal cells were considerably more permissive to DCV replication than glial or S2 cells, whereas independently evolved viral populations acquired distinct mutational patterns following propagation in mesodermal and S2 cells. These findings establish that cell type-specific intracellular environments can influence both viral fitness and evolutionary trajectories, even within the same host species.

Mesodermal cells consistently supported higher levels of viral RNA accumulation, VP1 production, and viral infectivity than either glial or S2 cells. Because S2 cells are derived from embryonic hemocytes, they are widely used as a model for antiviral immunity in *Drosophila*. Our transcriptomic and proteomic analyses further showed that S2 cells mounted stronger immune-associated responses than mesodermal cells, including enrichment of immune response and JAK–STAT signaling modules. Nevertheless, functional analyses demonstrated that the enhanced replication of DCV in mesodermal cells cannot be simply attributed to deficiencies in the three major antiviral pathways [7–11] examined. RNA interference was fully functional in all three cell types, the cGAS–STING pathway remained inducible by 2′3′-cGAMP, and JAK–STAT signaling exhibited cell type-dependent responsiveness without explaining the selective enhancement of DCV replication in mesodermal cells. Likewise, neither viral attachment and entry nor intracellular ATP abundance differed in a manner consistent with the observed differences in viral replication. Collectively, these observations suggest that additional cellular processes, rather than the canonical antiviral pathways examined here, determine the permissiveness of individual *Drosophila* cell types to DCV infection.

The molecular basis of this differential permissiveness remains unclear. Comparative transcriptomic and proteomic analyses identified numerous differences between mesodermal and S2 cells that extend beyond classical antiviral pathways. In particular, proteins involved in actin filament-based process, protein folding, and autophagy were differentially regulated following infection. Many of these pathways have previously been implicated in the replication of positive-strand RNA viruses, including dicistroviruses [21–25]. Rather than relying on a single antiviral pathway, DCV replication is therefore likely influenced by the combined effects of multiple host cellular processes, creating distinct intracellular selective environments in different cell types.

These differences in cellular environments were accompanied by distinct evolutionary trajectories during serial passage. Independent propagation of DCV in mesodermal and S2 cells altered viral infectivity toward the corresponding cell type, demonstrating rapid phenotypic adaptation during experimental evolution. However, these changes were not accompanied by detectable differences in viral RNA accumulation in adult flies, indicating that adaptation to a homogeneous cell-culture environment did not confer a measurable advantage during early infection of the whole-animal host. Similar discrepancies between cell culture adaptation and organismal infection have been reported for several RNA viruses [26–28], reflecting the substantially greater complexity of the multicellular host environment.

Genome sequencing further demonstrated that evolutionary constraints differed markedly across the DCV genome. The nonstructural proteins accumulated relatively few recurrent amino acid substitutions despite limited evidence of positive selection acting on the 2B gene in S2 cells. Furthermore, no parallel amino acid substitutions were identified in 1A, 2B, or 2C across independently evolved lineages. These observations indicate that the nonstructural proteins are subject to strong evolutionary constraint during short-term adaptation in cell culture.

By comparison, structural proteins exhibited broader signatures of positive selection, consistent with the greater accumulation of amino acid substitutions in ORF2. The most notable example was the VP3 G8089A mutation, which independently emerged in all three S2-passaged lineages but was absent from mesodermal-passaged viruses. Independent appearance of the same mutation under identical selective conditions is consistent with parallel evolution [29–31] and suggests that this genomic position may provide a selective advantage specifically in the S2 cellular environment. Nevertheless, the frequency of this mutation varied substantially among lineages, indicating that adaptation remains influenced by stochastic evolutionary processes during serial passage. Structural modeling predicted that the resulting R270H substitution is unlikely to substantially alter the interaction between VP3 and VP2, implying that any adaptive effect may instead involve more subtle changes in capsid stability, particle dynamics, receptor interactions, or other aspects of the viral life cycle. Experimental validation using reverse genetics will be required to determine the functional significance of this mutation.

Our study has several limitations. Viral populations were initially analyzed by pooled whole-genome sequencing to identify mutations accumulated during experimental evolution, whereas lineage-specific sequencing focused on candidate genomic regions. Consequently, additional low-frequency mutations outside these regions may have escaped detection in individual lineages. Furthermore, although serial passage provides a powerful approach for investigating viral adaptation, cultured cells represent simplified environments compared with intact tissues, where multiple cell types and systemic immune responses interact simultaneously. Thus, our study based on the cell culture adaptation could be different from the previous studies examining DCV evolution in whole animals [32–35].

In conclusion, our results demonstrate that different *Drosophila* cell types provide distinct intracellular environments that shape both DCV replication and viral evolution. Rather than being determined solely by the major antiviral pathways examined here, viral adaptation appears to emerge from the integrated effects of numerous host cellular processes that collectively define the intracellular selective landscape. These findings provide a framework for understanding how tissue-specific cellular environments contribute to RNA virus evolution and highlight the importance of considering host cell identity when investigating virus–host interactions and viral adaptation.

## Methods

### Virus stock and titration

The DCV stock, kindly provided by Dr. Tao Peng (Guangzhou Medical University), was propagated in *Drosophila* S2 cells cultured at 25 °C in Schneider’s medium supplemented with 10 % fetal bovine serum, 50 U/mL penicillin, and 50 μg/mL streptomycin (Beyotime).

Viral titers were determined by an endpoint dilution assay. S2 or mesodermal cells were seeded in flat-bottom 96-well plates at 3 × 10^4^ cells per well in 75 μL of culture medium. The following day, 10-fold serial dilutions of the virus stock were prepared in round-bottom 96-well plates containing 180 μL of culture medium per well by adding 20 μL of virus suspension to the first well and sequentially transferring 20 μL through 12 dilutions. Aliquots of each dilution (25 μL) were added to four replicate wells containing cells. After incubation at 25 °C for 2 days, the cells were fixed with 100 μL of 8 % paraformaldehyde in PBS for 20 min, permeabilized with PT buffer (PBS containing 0.1 % Triton X-100) for 5 min, and blocked with PT containing 5 % bovine serum albumin for 30 min. The cells were incubated overnight at 4 °C with rabbit anti-DCV VP1 antibody diluted 1:3,000 in blocking solution. After three 10-min washes with PT, the cells were incubated for 3 h at 25 °C with horseradish peroxidase-conjugated anti-rabbit IgG (Proteintech; 1:3,000) diluted in PT containing 5 % normal goat serum. Following three additional washes with PT, VP1-positive cells were visualized using the AEC Chromogen Kit (red; Boster). Wells containing more than 10 VP1-positive cells were scored as positive. Viral titers were expressed as the median tissue culture infectious dose (TCID50) and calculated using the Reed–Muench method.

### Transcriptomic and proteomic analyses of DCV-infected glial, mesodermal, and S2 cells

Glial, mesodermal, and S2 cells were seeded in 10-cm culture dishes at 1 × 10^7^ cells per dish and infected the following day with DCV at 3 × 10^7^ TCID50. Cells were collected at 6, 12, and 24 h post-infection, with three biological replicates per condition. Uninfected control cells were prepared in parallel. Cells were washed twice with PBS and lysed in 1 mL TRIzol reagent for total RNA extraction. Total RNA (20 μg) from each sample was submitted to the Beijing Genomics Institute (BGI; Shenzhen, China) for RNA sequencing on the DNBSEQ-T7 platform, generating at least 6 GB of clean data per sample. Raw reads were processed using SOAPnuke v2.2.1 [36] to remove low-quality, adapter-contaminated, and reads containing more than 5 % ambiguous bases, yielding clean reads with Q20 ≥ 90 % and Q30 ≥ 85 %. Clean reads were aligned to the *D. melanogaster* reference genome (GCF_000001215.4_Release_6_plus_ISO1_MT) using HISAT2 v2.2.1 [37] with the strand-specific parameter --rna-strandness RF. Transcripts were assembled using StringTie v2.2.1 [38], and assemblies from all samples were merged using Cuffmerge v2.2.1 [39] to generate a comprehensive transcriptome reference. Gene- and transcript-level expression was quantified using RSEM v1.3.1 [40], with Bowtie2 v2.4.5 [41] as the alignment engine, and expression values were reported as fragments per kilobase of transcript per million mapped reads (FPKM). The --keep-intermediate-files option was used to retain unmapped reads for subsequent viral RNA analysis. Principal component analysis (PCA) was performed on the merged gene-level transcripts per million (TPM) matrix using the fast.prcomp function in the gmodels package in R v4.4. PCA plots were generated using ggpubr and ggplot2, with samples grouped by cell type and infection time.

For proteomic analysis, mesodermal and glial cells were washed with PBS for 5 min and treated with 1 mL of 0.5 % trypsin (Beyotime) for 10 min at 25 °C. Cells were resuspended in 3 mL serum-free Schneider’s medium and collected by centrifugation. Cell pellets were resuspended in 300 μL of 8 M urea and sonicated, and the soluble protein fractions were collected by centrifugation. Protein samples (40 μL, containing 20–223 μg protein) were submitted to BGI for label-free quantitative proteomic analysis using data-independent acquisition (DIA). Peptides and proteins were identified and quantified by deconvolution of the DIA data against a data-dependent acquisition spectral library. Mass spectrometry data from 36 samples were acquired in DIA mode using a Q Exactive HFX mass spectrometer (Thermo Fisher Scientific), resulting in the quantification of 89,506 peptides and 6,303 proteins. Protein intensities were summarized and normalized by median equalization using MSstats [42]. PCA was performed using the normalized abundance values of all quantified proteins, and the first two principal components were examined to assess biological replicate reproducibility and differences among experimental groups.

### Quantification of DCV genomic RNA in the transcriptome

The complete DCV reference genome (NC_001834.1) and corresponding GFF annotation file were downloaded from the NCBI Nucleotide database. A Bowtie2 index was generated using bowtie2-build, and the GFF file was converted to GTF format using gffread v0.11.2 [43] for gene-length calculation. To quantify viral RNA, reads that remained unmapped after RSEM analysis were aligned to the DCV reference genome using Bowtie2 in local alignment mode with the --sensitive preset. SAM files were converted to sorted BAM files using SAMtools v1.9 [44], and genome-level read counts were obtained using samtools idxstats. Relative DCV genome abundance was estimated as the number of mapped reads multiplied by the read length and divided by the viral genome length. Samples containing detectable DCV reads were retained for gene-level quantification.

To quantify reads at individual viral genes (ORF1 and ORF2), we implemented a custom R-based counting algorithm following the viGEN methodology [45]. For each sample, the sorted BAM file was subsetted to the DCV reference using the Rsamtools package (v2.8.0). Read positions were extracted from alignments, and for each annotated feature (gene or CDS) from the DCV GFF file, reads were categorized as: (1) fully contained within the feature (in-region), (2) overlapping feature boundaries (on-boundary), or (3) falling outside any annotated feature (in-gaps). The final gene-level count was defined as the sum of in-region and on-boundary counts. This count matrix was collated across all 36 samples.

For joint analysis of host and viral transcripts, RSEM-derived expected counts for host genes were combined with DCV gene counts into a single count matrix. Gene lengths used for transcripts-per-million (TPM) normalization were calculated from non-overlapping exonic regions using the GenomicFeatures package v1.46.0 [46]. TPM values were calculated using effective gene length and mean fragment length as described by Wagner et al. [47]. Genes with an effective length shorter than the mean fragment length were excluded from the normalization.

### Integration of omics datasets and definition of functional modules

Differential expression of host transcripts was analyzed using DESeq2 [48]. For each cell type (glial, mesodermal, and S2), DCV-infected samples collected at 6, 12, and 24 h post-infection were compared with the corresponding mock-infected control samples. Genes with an absolute log2fold change >1 and an unadjusted *P-*value <0.001 at any time point were defined as infection-regulated transcripts. Differential protein abundance was analyzed using MSstats for the same cell type- and time point-specific comparisons. Proteins with an absolute log2fold change >1 and a Benjamini–Hochberg-adjusted *P-*value <0.05 at any time point were considered infection-regulated. Protein identifiers were converted to FlyBase gene identifiers using the UniProt ID mapping service. For each cell type, transcriptomics- and proteomics-derived infection-regulated genes were combined to generate an integrated gene set for network-based functional analysis.

A high-confidence *D. melanogaster* protein–protein interaction network was obtained from STRING v12.0 [49] and restricted to proteins represented in each integrated gene set using the STRING gene alias table. Only interactions with a combined confidence score >600 were retained. Functionally related modules were identified using the network-based clustering framework described by Bouhaddou et al. [50]. Diffusion State Distance (DSD), which measures similarity between random-walk diffusion profiles, was calculated for all gene pairs in the extracted network [51]. The resulting distance matrix was subjected to average-linkage hierarchical clustering, and modules were defined using the cutreeHybrid function in the dynamicTreeCut R package v1.63-1 [52], with deepSplit = 3. This procedure was performed separately for the glial, mesodermal, and S2 cell gene sets.

Modules were annotated by Gene Ontology enrichment analysis using clusterProfiler v4.4.1 [53] and the *D. melanogaster* OrgDb resource AH107058, obtained through AnnotationHub v4.2.2. For each module, the eight GO terms with the lowest *P-*values were retained. The term containing the largest number of genes from the module was used as the module name; ties were resolved by selecting the term with the lower *P-*value.

### Generation of anti-DCV VP1 antibody

The C-terminal region of DCV VP1 (amino acids 167–265) was amplified by PCR using the primer pair DCV752-F-BamHI and DCV850-R-NotI. The PCR product was digested with BamHI and NotI, purified, and ligated into the corresponding restriction sites of the pGEX-6P-3 expression vector (Cytiva). The resulting plasmid was transformed into *Escherichia coli* BL21 cells. Transformed *E. coli* BL21 cells were cultured in 1 L of LB medium containing 1 % glucose and 0.1 mg/mL ampicillin at 37 °C until the optical density at 600 nm reached approximately 0.5. The culture was then cooled on ice, and recombinant protein expression was induced with 0.1 mM isopropyl β-D-1-thiogalactopyranoside at 15 °C for 16 h.

Bacterial cells were harvested by centrifugation and resuspended in 100 mL of ice-cold TNED buffer (50 mM <u>T</u>ris-HCl, pH 8.0, 150 mM <u>N</u>aCl, 2 mM <u>E</u>DTA, and 1 mM <u>D</u>TT) supplemented with 0.5 % Triton X-100 and a protease inhibitor cocktail (Beyotime). Cell lysates were prepared by sonication using a Q700 Sonicator (Qsonica) at an amplitude of 100 on ice for 45 min (30-s pulses with 3-min intervals). After centrifugation, the supernatant was incubated with 1 mL of BeyoGold™ GST-tag Purification Resin (Beyotime) at 4 °C for 2 h with gentle rotation. The resin was subsequently washed five times with 10 mL of TNED buffer. To release the recombinant VP1 protein, 2 mL of TNED buffer containing 100 units of PreScission Protease (Beyotime) was added to the resin, followed by incubation overnight at 4 °C. The supernatant containing the cleaved VP1 protein was collected, and the resin was further eluted once with 2 mL of TNED buffer. The eluate was combined with the initial supernatant and dialyzed twice against 2 L of PBS containing 1 mM EDTA and 0.5 mM DTT at 4 °C for 24 h. The purified VP1 protein was then sent to GeneScript (Nanjing, China) for the production of rabbit polyclonal anti-DCV VP1 antibody.

### Quantification of DCV genomic RNA and VP1 protein in infected cells

Glial, mesodermal, and S2 cells were seeded in 24-well plates at 2.5 × 10^5^ cells per well, using five biological replicates for DCV genomic RNA (gRNA) quantification and three for VP1 protein quantification. Cells were infected with 1,000 TCID50 of DCV and harvested at 24 h post-infection. For gRNA quantification, 0.2 μg of total RNA was reverse-transcribed in a 20 μL reaction using ReverTra Ace (TOYOBO) and random primers. The resulting cDNA was diluted twofold with water and analyzed by qPCR using the primers DCV-qPCR-F and DCV-qPCR-R. *D. melanogaster* 18S rRNA, amplified using Dm18S-F and Dm18S-R, served as the reference transcript. Relative DCV gRNA abundance was calculated using the 2^−ΔΔCt^ method, with one infected S2 cell sample used as the calibrator and assigned a value of 1. Statistical significance was assessed using the Kruskal–Wallis test followed by one-sided Steel multiple-comparison tests.

For VP1 quantification, cells from each well were lysed in 100 μL of RIPA buffer containing 20 mM Tris–HCl, pH 7.5, 150 mM NaCl, 1 % NP-40, 0.5 % sodium deoxycholate, 0.1 % SDS, and a protease inhibitor cocktail (Beyotime). Lysates were clarified by centrifugation, and protein concentrations were determined using the BCA Protein Assay Kit (Beyotime). Equal amounts of protein (10 μg) were separated on duplicate 10 % SDS–PAGE gels. One gel was stained with InstantBlue (Abcam) to assess protein loading, whereas proteins in the second gel were transferred to a nitrocellulose membrane (Pall Life Sciences). The membrane was blocked with 5 % BSA in PBST (PBS containing 0.1 % Tween-20) and incubated overnight at 4 °C with rabbit anti-DCV VP1 antibody diluted 1:1,000. After five 5 min washes with PBST, the membrane was incubated for 2 h at room temperature with IRDye 680RD donkey anti-rabbit IgG (H+L) secondary antibody (LI-COR Biosciences; 1:10,000) diluted in PBST containing 5 % skim milk. Following additional washes, fluorescence was detected using a ChemiDoc MP Imaging System (Bio-Rad). VP1 band intensities were quantified after background subtraction using ImageJ. Statistical significance was assessed by one-way ANOVA followed by one-sided Dunnett’s multiple-comparison tests.

### Comparison of RNAi activity in glial, mesodermal, and S2 cells

Glial, mesodermal, and S2 cells were seeded in 6-well plates at 1 × 10^6^ cells per well and co-transfected with 2 μg of a GFP-expressing pAc5.1/V5-His B plasmid (Thermo Fisher Scientific) and 2 μg of either GFP-specific or mCherry-specific control dsRNA using HilyMax (Dojindo). Cells were analyzed 2 days after transfection. Templates for dsRNA synthesis were amplified by PCR using the primer pairs T7-GFP-F/T7-GFP-R or T7-mCherry-F/T7-mCherry-R and the GFP- or mCherry-expressing plasmid (Addgene plasmid #55045), respectively. RNA was synthesized using T7 RNA polymerase (Takara), and complementary RNA strands were annealed to generate dsRNA. The sizes of the PCR products and dsRNAs were verified by agarose gel electrophoresis.

GFP fluorescence was examined using a Nikon Eclipse Ti2-U fluorescence microscope with the exposure settings indicated in the figure legend. GFP expression was also quantified using a four-color BD FACSCelesta flow cytometer (BD Biosciences). Data were acquired using the automatic acquisition and compensation settings, with FL1–FL4 photomultiplier tube voltages adjusted manually, and analyzed using FlowJo. For immunoblotting, 2 × 10^5^ cells were lysed in 100 μL of SDS–PAGE sample buffer containing 2 % SDS, 10 % glycerol, 10 % β-mercaptoethanol, 0.25 % bromophenol blue, and 50 mM Tris–HCl, pH 6.8. Lysates were heated at 95 °C for 5 min, and 25 μL of each sample was separated on duplicate 10 % SDS–PAGE gels. One gel was stained with InstantBlue, whereas proteins from the other were transferred to a nitrocellulose membrane and immunoblotted as described above using an anti-GFP polyclonal antibody (Proteintech; 1:1,000).

### Comparison of cGLR–STING signaling in glial, mesodermal, and S2 cells

Glial, mesodermal, and S2 cells were seeded in 96-well plates at 4 × 10^4^ cells per well across 12 wells per cell type. Individual wells were transfected with 0.8 μg of 2′3′-cGAMP or an equivalent volume of water as the control using 2 μL of HilyMax and incubated for 8 h. Total RNA was extracted from each well and dissolved in 20 μL of water, of which 4 μL was used for reverse transcription as described above. The resulting cDNA was diluted threefold, and 1 μL was used for qPCR analysis of *Srg3* mRNA and *D. melanogaster* 18S rRNA using the primer pairs DmSrg3-F/DmSrg3-R and Dm18S-F/Dm18S-R, respectively. Relative *Srg3* expression was calculated using the 2^−ΔΔCt^ method, with one untreated S2 cell sample used as the calibrator and assigned a value of 1. Differences between cGAMP-treated and control samples within each cell type were analyzed using two-sided Welch’s *t*-tests. Comparisons among the three cell types were performed using the Kruskal–Wallis test followed by one-sided Steel multiple-comparison tests. To quantify endogenous *cGLR1* expression, glial, mesodermal, and S2 cells were seeded in 24-well plates at 2.5 × 10^5^ cells per well, with five replicate wells per cell type. Total RNA was extracted the following day, and *cGLR1* mRNA was quantified by RT–qPCR using the primers DmcGLR1-F and DmcGLR1-R. Differences among the three cell types were analyzed using the Kruskal–Wallis test followed by one-sided Steel multiple-comparison tests.

### Comparison of JAK–STAT signaling in mesodermal, glial, and S2 cells

To generate the Upd2 expression construct, the *Upd2* open reading frame was amplified from S2 cell cDNA using the primers Upd2-5-EcoRI and Upd2-3-XhoI. The PCR product was digested with EcoRI and XhoI and inserted into the corresponding sites of pAc5.1/V5-His B. Mesodermal, glial, and S2 cells were seeded in 96-well plates at 4 × 10^4^ cells per well, with 10 replicate wells per cell type, and transfected with 0.2 μg of either the Upd2 expression plasmid or empty pAc5.1/V5-His B as a mock control using 1 μL HilyMax. After 2 days, *TotM* mRNA and *D. melanogaster* 18S rRNA were quantified by RT–qPCR using the primer pairs DmTotM-F/DmTotM-R and Dm18S-F/Dm18S-R, respectively. Relative *TotM* expression was calculated using the 2^−ΔΔCt^ method, with one mock-transfected S2 cell sample used as the calibrator and assigned a value of 1. Differences between Upd2-expressing and mock-transfected cells within each cell type were analyzed using two-sided Welch’s *t*-tests, whereas comparisons among the three cell types were performed using the Kruskal–Wallis test followed by the Steel– Dwass test. Endogenous *Upd2* expression was quantified using the cDNA samples prepared for *cGLR1* analysis, whereas ectopic *Upd2* expression was measured using cDNA from the transfected cells. In both cases, *Upd2* mRNA was quantified using the primers DmUpd2-F and DmUpd2-R, and differences among cell types were analyzed using the Kruskal–Wallis test followed by the Steel– Dwass test.

### Assays of DCV binding and entry

DCV binding and entry were assessed in mesodermal and S2 cells seeded in 24-well plates at 2.5 × 10^5^ cells per well, with five replicate wells per condition. For the binding assay, cells were preincubated at 4 °C for 15 min, after which the medium was replaced with 100 μL of ice-cold medium containing 4 × 10^6^ TCID50 of DCV. Following incubation at 4 °C for 30 min, cells were washed twice with 200 μL of ice-cold PBS containing 1 mM MgCl2 and 1 mM CaCl2, and lysed in 200 μL of TRIzol reagent for total RNA extraction.

For the entry assay, cells were incubated with 100 μL of medium containing 4 × 10^6^ TCID50 of DCV at 25 °C for 2 h. Cells were then washed twice with PBS containing 1 mM MgCl2 and 1 mM CaCl2 at room temperature. Surface-bound virus was removed by treatment with 0.5 % trypsin for 10 min at room temperature, followed by two washes with 200 μL of ice-cold culture medium. Cells were lysed in 200 μL of TRIzol reagent, and DCV genomic RNA and *D. melanogaster* 18S rRNA were quantified by RT–qPCR as described above. Relative DCV genomic RNA abundance was calculated using the 2^−ΔΔCt^ method, with one S2 cell sample from the binding assay used as the calibrator and assigned a value of 1. Differences between mesodermal and S2 cells were analyzed using two-sided Welch’s *t*-tests.

### Measurement of intracellular ATP concentration

Mesodermal and S2 cells were seeded in opaque 96-well plates at 3 × 10^4^ cells per well, with five replicate wells per cell type. The following day, the plates were equilibrated at room temperature for 10 min, after which 100 μL of CellTiter-Lumi™ reagent (Beyotime) was added to each well. The plates were shaken at room temperature for 2 min to lyse the cells and incubated for an additional 10 min to stabilize the luminescence signal. Luminescence was measured using a Varioskan™ LUX multimode microplate reader (Thermo Fisher Scientific), and ATP content was calculated from an ATP standard curve. Intracellular ATP concentrations were estimated by normalizing ATP content to the estimated total cellular volume. Individual cell volumes were calculated from the mean cell diameters (mesodermal cells, 8.87 ± 1.28 μm; S2 cells, 9.20 ± 1.71 μm), assuming a spherical cell shape.

### Experimental evolution of DCV

The original DCV stock was subjected to three sequential bottlenecks in S2 cells by inoculating cells in 96-well plates with the highest viral dilution that produced a cytopathic effect. The resulting bottlenecked virus was expanded in S2 cells cultured in 6-well plates for 5 days. Culture supernatants were collected, aliquoted, stored at −80 °C, and titrated; this preparation was designated the parental DCV stock. For the first passage, 2 × 10^6^ mesodermal or S2 cells seeded in 6-cm dishes were infected with 10,000 TCID50 of parental DCV and incubated for 4 days. Five independent viral lineages were established for each cell type from passage 1 onward. At each subsequent passage, cells were infected with 10,000 TCID50 of the corresponding evolved virus. After 4 days, culture supernatants were collected, aliquoted, stored at −80 °C, and titrated for use in the next passage.

### Infection of cultured cells and adult flies with parental and evolved DCV

To compare early replication of parental and evolved DCV, equal volumes of culture supernatant from the five P15 lineages of each cell type were pooled. DCV genomic RNA (gRNA) concentrations in the parental, mesodermal P15, and S2 P15 stocks were determined by RT–qPCR using a standard curve generated from a DCV PCR product. Because the gRNA concentration of the parental stock was approximately half that of either P15 stock, viral inputs were normalized by dilution. Mesodermal and S2 cells were seeded in 96-well plates at 4 × 10^4^ cells per well and infected with 1 μL of parental DCV diluted 1:50 or mesodermal or S2 P15 DCV diluted 1:100. After 24 h, total RNA was extracted and DCV gRNA was quantified by RT–qPCR. Relative gRNA abundance was calculated using the 2^−ΔΔCt^ method, with one parental DCV-infected S2 cell sample used as the calibrator and assigned a value of 1. Mesodermal and S2 cells infected with the same viral stock were compared using two-sided Welch’s *t*-tests. The three viral stocks were compared within each cell type using the Kruskal–Wallis test followed by Steel–Dwass multiple-comparison tests. For VP1 quantification, infected cells were lysed in 70 μL of SDS–PAGE sample buffer, and 10 μL of each sample was separated on duplicate 10 % SDS–PAGE gels. One gel was stained with InstantBlue, whereas the other was used for immunoblotting with anti-DCV VP1 antibody as described above. VP1 abundance between mesodermal and S2 cells infected with the same stock was compared using two-sided Welch’s *t*-tests, whereas the three viral stocks were compared within each cell type using one-way ANOVA followed by the Tukey–Kramer test.

For infection of adult flies, 3–5 day old female *w*^1118^ flies (Bloomington Drosophila Stock Center #3605) were injected intrathoracically with 50 nL of parental DCV diluted 1:50 or mesodermal or S2 P15 DCV diluted 1:100 in 10 mM Tris–HCl, pH 7.3, using a Nanoject II microinjector (Drummond Scientific). At 24 h post-infection, five biological replicates, each consisting of a pool of ten flies, were collected for RT–qPCR analysis. DCV gRNA levels among the three viral stocks were compared using one-way ANOVA.

### NGS library preparation

Total RNA was extracted from 200 μL of each virus stock using 1 mL of TRIzol reagent, with 10 μg of glycogen added as a carrier, and dissolved in 20 μL of water. After reverse transcription of the 5 μL RNA sample, viral cDNA was amplified using KOD FX Neo DNA polymerase (TOYOBO) as three overlapping amplicons of approximately 3.2 kb using the primer pairs 26F/3226R, 3022F/6223R, and 5988F/9102R for fragments 1, 2, and 3, respectively. The amplicons were fragmented using a Bioruptor Pico sonicator (Diagenode) in 1.5-mL Bioruptor tubes containing 0.1 % SDS for 30 cycles of 30s sonication followed by 30s cooling. Fragmented DNA was purified using a QIAquick PCR Purification Kit (QIAGEN), and 1 μg was used for library preparation with the NEBNext Ultra II DNA Library Prep Kit (E7645; New England Biolabs). Libraries were indexed using NEBNext Multiplex Oligos (E7600; New England Biolabs) and sequenced on an Illumina NextSeqPE300 platform using paired-end 100 bp reads. Base calling was performed using Illumina Real-Time Analysis software, and reads were demultiplexed according to their index sequences using bcl2fastq v2.20. Read numbers and the proportion of bases with quality scores ≥Q30 were recorded. All primers used in this study are listed in Supplementary Data 2.

### NGS data processing

The DCV EB strain genome (GenBank accession NC_001834.1) was used as the initial reference. Raw reads from the parental viral stock were processed using a custom workflow incorporating V-pipe [54] for quality control and preprocessing. Reads were aligned to the EB reference using BWA-MEM v0.7.18 [55] and sorted using SAMtools v1.21 [44]. Variants were called using the mpileup and call functions in bcftools v1.21 [56], and the parental consensus sequence was generated using bcftools consensus. For fragment-specific analyses, the parental consensus was restricted to the corresponding genomic regions: BN1, positions 26–3226; BN3, positions 5988–9102; and BN1-3, the full-length consensus sequence. A 1-bp deletion at position 329 in the parental stock relative to the EB reference was incorporated into subsequent coordinate conversions.

Raw paired-end reads were filtered using PRINSEQ-lite v0.20.4 [57] with the parameters -ns_max_n 4 -min_qual_mean 20 -trim_qual_left 20 - trim_qual_right 20 -trim_qual_window 10, and read quality was assessed using FastQC [58]. For each dataset (BN1, BN3, and BN1-3), filtered reads were aligned to the corresponding parental consensus sequence using BWA-MEM v0.7.18 and sorted using SAMtools v1.21.

SNVs and viral haplotypes were reconstructed using ShoRAH v1.99.2 [59] with default parameters, including a posterior probability threshold of 0.9, α = 0.1, and a shift value of 3. Variants were filtered using the ShoRAH strand-bias test [60] and were required to meet a Phred quality threshold corresponding to an estimated error rate of 0.0001. Mutation coordinates were converted to the EB reference coordinate system according to the genomic position of each amplicon. Variants were annotated using an adapted version of vcf-annotator. Standard annotation was performed using the GenBank reference sequence, whereas corrected annotation used the REF allele in each VCF file to reconstruct the reference codon, thereby accounting for differences between the parental and GenBank sequences, including indels.

Population nucleotide diversity (π), Shannon entropy, and the number of polymorphic sites were calculated using custom Python scripts in Python v3.11.12, with a minimum variant-frequency threshold of 0.0001. Co-occurring mutations were identified from ShoRAH-reconstructed haplotypes. Nucleotide diversity at synonymous and nonsynonymous sites (πS and πN, respectively) and πN-πS was calculated using SNPGenie v2019.10.31 [61] in within-pool mode with the same minimum-frequency threshold. Base-quality and coverage statistics were obtained using pysamstats.

### Alignment of ORF2 amino acid sequences of dicistroviruses

ORF2 amino acid sequences from seven dicistroviruses, including DCV and Cricket paralysis virus (CrPV), were retrieved from the NCBI database and aligned using Clustal Omega with default parameters.

### AlphaFold3 prediction of the DCV VP1–VP4 complex

The individual amino acid sequences of DCV VP1, VP2, VP3, and VP4 were submitted to the AlphaFold3 server to predict the structure of the VP1–VP4 complex. The predicted local distance difference test (pLDDT) scores exceeded 70 for most residues, except near the C-terminal region of VP2, indicating generally high confidence in the predicted structure. Structural visualization and analysis were performed using UCSF ChimeraX v1.12 [62].

## Data Availability

The datasets generated and analyzed during the current study have been deposited in publicly available repositories. The RNA-seq raw data have been submitted to the NCBI Sequence Read Archive (SRA) under BioProject accession [PRJNA1496022]. The mass spectrometry proteomics data have been deposited to the ProteomeXchange Consortium via the PRIDE [63] partner repository with the dataset identifier [PXD081430]. All raw sequencing data have been deposited in NCBI BioProject under accession number [PRJNA1503189].

## Acknowledgements

This work was supported by Jinji Lake Double Hundred Talents Programme to TK. The funder had no role in study design, data collection and analysis, decision to publish, or preparation of the manuscript.

## Authorship contributions

Tatsuhiko Kadowaki conceived and designed research strategy and wrote the paper. Qinyi Liang, Jiaxin Liu, Yunfun Huang, and Xuye Yuan performed the experiments.

## Competing interests

The authors declare no competing interests.

## Supplementary information

Supplementary Data 1

Functional pathway modules altered by DCV infection Supplementary Data 2

List of primers used in this study.

Supplementary Figure 1

VP1 accumulation in mesodermal and S2 cells infected with parental or evolved DCV

Supplementary Figure 2

Mutation dynamics and selection in the DCV ORF1 region encoding 1A, 2B, and 2C

Supplementary Figure 3

Mutation dynamics and selection in the DCV ORF2 region encoding structural proteins

---

**Supplementary Figure 1 VP1 accumulation in mesodermal and S2 cells infected with parental or evolved DCV**

(A) Immunoblot analysis of DCV VP1 in mesodermal and S2 cells at 24 h after infection with parental DCV or pooled mesodermal P15 or S2 P15 viruses. Viral inputs were normalized according to DCV genomic RNA abundance. Total protein visualized by InstantBlue staining served as the loading control. The VP1 band is indicated by an arrowhead, and molecular mass markers are shown in kilodaltons (kDa) on the left. (B) Quantification of VP1 band intensities in panel A. Mesodermal and S2 cells infected with the same viral stock were compared using two-sided Welch’s *t*-tests. The three viral stocks were compared within each cell type using one-way ANOVA followed by the Tukey–Kramer test. Horizontal lines indicate means, error bars indicate standard deviations, and symbols represent biological replicates; n = 3.

**Supplementary Figure 2 Mutation dynamics and selection in the DCV ORF1 region encoding 1A, 2B, and 2C**

(A) Number of SNVs detected in mesodermal- and S2-passaged DCV populations at P5, P10, and P15. SNV numbers differed between the two cell types only at P5 (Mann–Whitney *U*-test). (B) Distribution of SNVs among four frequency ranges: 0.01–0.1 %, 0.1–0.5 %, 0.5–50 %, and >50 %. No significant differences were detected between mesodermal- and S2-passaged populations in any frequency range (two-sided Welch’s *t*-test). (C) Mean nucleotide diversity of mesodermal- and S2-passaged DCV populations at P5, P10, and P15. Nucleotide diversity was higher in S2-than in mesodermal-passaged populations at P10 (Mann–Whitney *U*-test). (D) Nucleotide diversity of the regions encoding 2B and 2C at each passage. Blue and orange symbols represent mesodermal- and S2-passaged populations, respectively, and crosses indicate the mean nucleotide diversity of the entire analyzed ORF1 region. Asterisks indicate significant differences from the corresponding regional mean (*P* < 0.05, one-sample *t*-test). (E) Nucleotide diversity at nonsynonymous (πN; red) and synonymous (πS; blue) sites in 1A, 2B, and 2C at each passage. Light-red and light-blue backgrounds indicate πN − πS > 0 and πN − πS < 0, respectively. Data represent the mean ± SD of three independently evolved viral lineages. ns, not significant.

**Supplementary Figure 3 Mutation dynamics and selection in the DCV ORF2 region encoding structural proteins**

(A) Number of SNVs detected in ORF2 of mesodermal- and S2-passaged DCV populations at P5, P10, and P15. The number of SNVs increased significantly in mesodermal-passaged populations between P5 or P10 and P15 (one-way ANOVA followed by the Tukey–Kramer test). (B) Distribution of SNVs among three frequency ranges: 0.1–0.5 %, 0.5–50 %, and >50 %. Mesodermal-passaged populations contained a higher proportion of intermediate-frequency SNVs (0.5–50 %), whereas S2-passaged populations contained a higher proportion of high-frequency SNVs (>50 %) (two-sided Welch’s *t*-test). (C) Mean nucleotide diversity of ORF2 in mesodermal- and S2-passaged populations at P5, P10, and P15. No significant change was detected across passages in either cell type. (D) Nucleotide diversity of the regions encoding VP2, VP3, and VP1 at each passage. Blue and orange symbols represent mesodermal- and S2-passaged populations, respectively, and crosses indicate the mean nucleotide diversity of the entire ORF2 region. Asterisks indicate significant differences from the corresponding ORF2 mean (*P* < 0.05, one-sample *t*-test). (E) Nucleotide diversity at nonsynonymous (πN; red) and synonymous (πS; blue) sites in VP2, VP4, VP3, and VP1 at each passage. Light-red and light-blue backgrounds indicate πN − πS > 0 and πN − πS < 0, respectively. Data represent the mean ± SD of three independently evolved viral lineages. ns, not significant.

## Notes

### Competing Interest Statement

The authors have declared no competing interest.

