## Supplementary figures and images for "Host cell identity shapes the replication and evolution of *Drosophila* C virus"

### Supplementary Figure 1

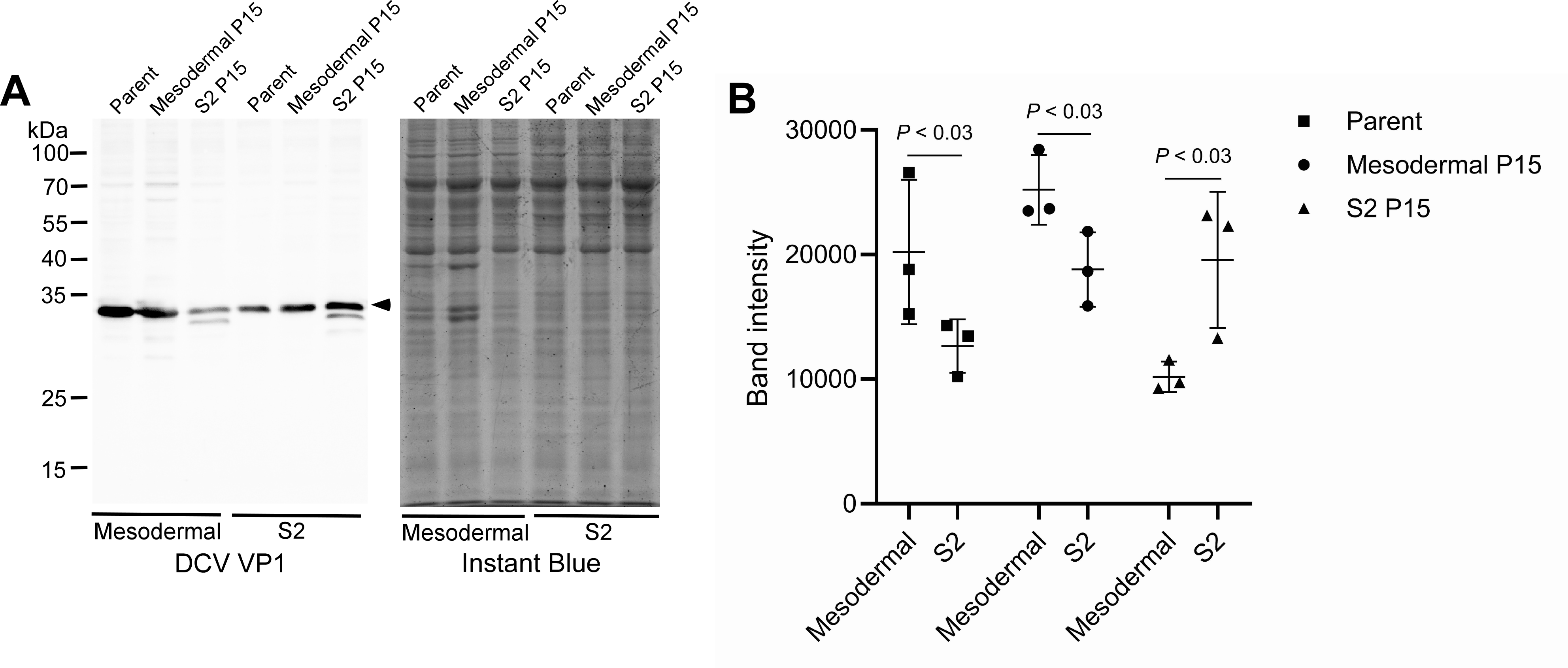

### Supplementary Figure 2

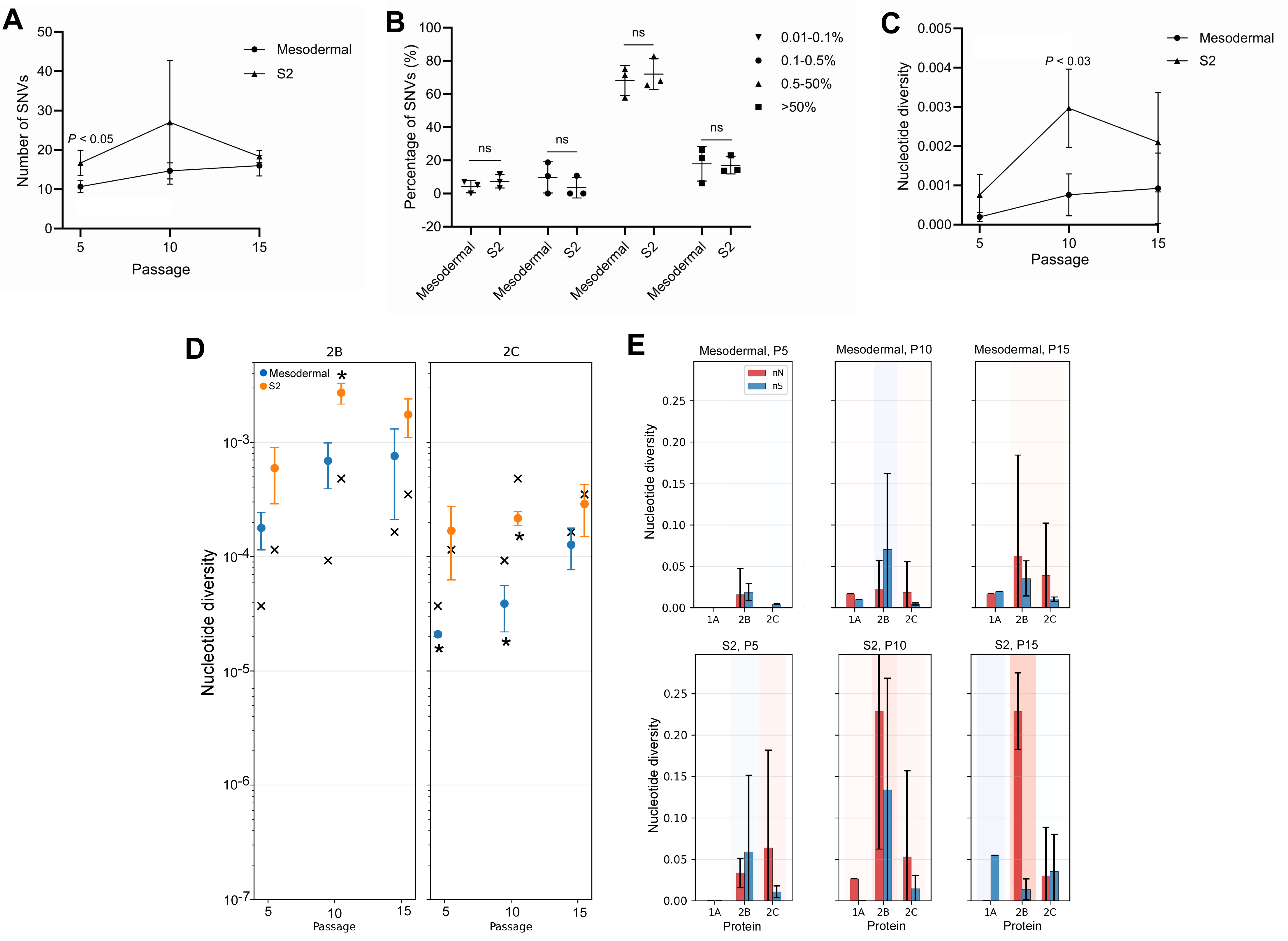

### Supplementary Figure 3

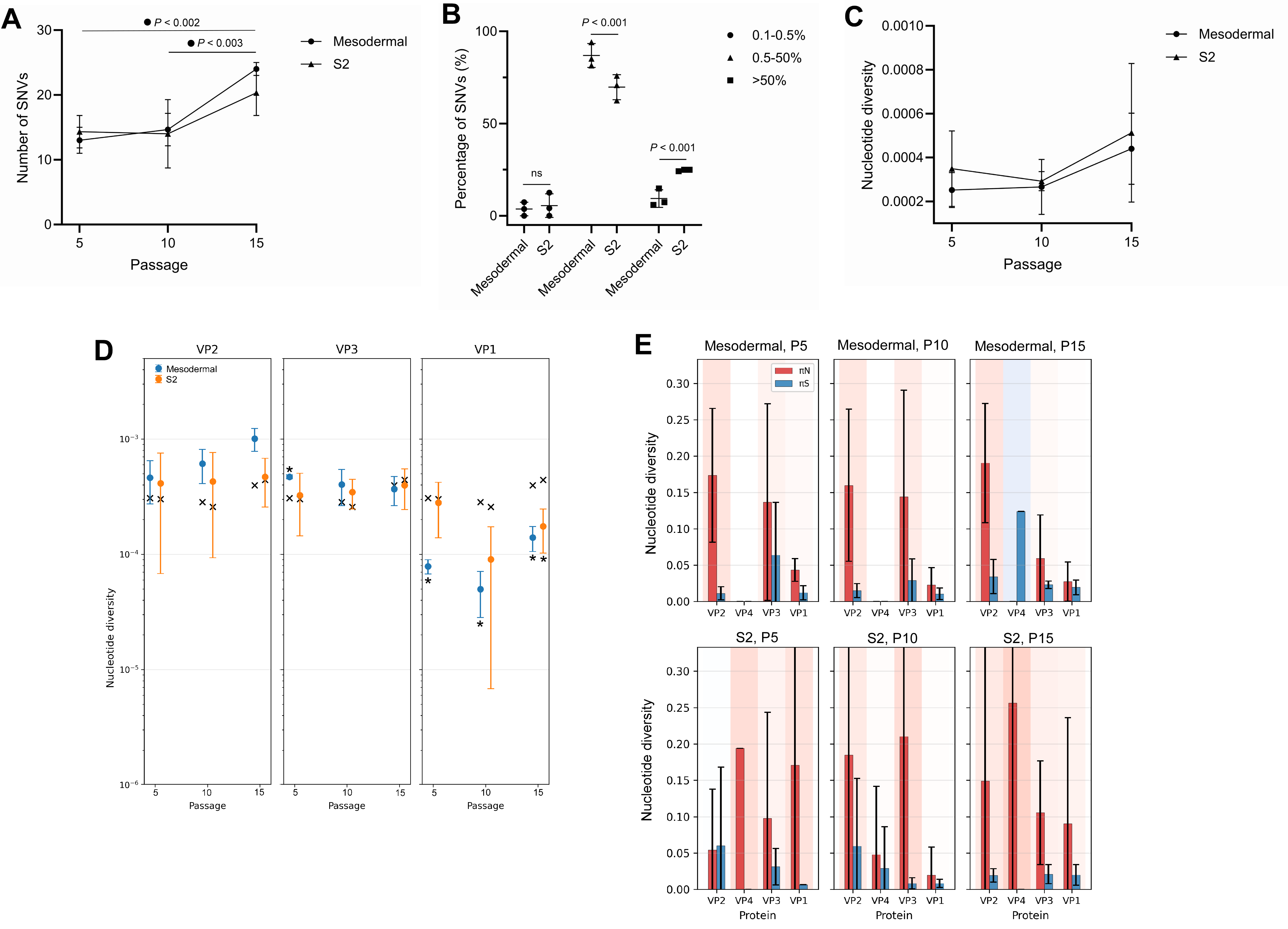
